# A generalizable experimental framework for automated cell growth and laboratory evolution

**DOI:** 10.1101/280867

**Authors:** Brandon G. Wong, Christopher P. Mancuso, Szilvia Kiriakov, Caleb J. Bashor, Ahmad S. Khalil

**Author notes:** Co-first author (Brandon G. Wong, Christopher P. Mancuso). Correspondence (C.J.B), (A.S.K.).

## Abstract

In the post-genomics era, exploration of phenotypic adaptation is limited by our ability to experimentally control selection conditions, including multi-variable and dynamic pressure regimes. While automated cell culture systems offer real-time monitoring and fine control over liquid cultures, they are difficult to scale to high-throughput, or require cumbersome redesign to meet diverse experimental requirements. Here we describe eVOLVER, a multipurpose, scalable DIY framework that can be easily configured to conduct a wide variety of growth fitness experiments at scale and cost. We demonstrate eVOLVER’s versatility by configuring it for diverse growth and selection experiments that would be otherwise challenging for other systems. We conduct high-throughput evolution of yeast across different population density niches. We perform growth selection on a yeast knockout library under temporally varying temperature regimes. Finally, inspired by large-scale integration in electronics and microfluidics, we develop novel millifluidic multiplexing modules that enable complex fluidic routines including multiplexed media routing, cleaning, vial-to-vial transfers, and automated yeast mating. We propose eVOLVER to be a versatile design framework in which to study, characterize, and evolve biological systems.

## INTRODUCTION

Biological organisms are embedded in complex, dynamically changing environments that shape their evolved phenotype^1–3^ (**Fig. 1a**). An integrated understanding of how phenotype is encoded and adapted to specific selective pressures remains a central challenge for biological research. In the laboratory, experiments involving growth selection in liquid culture have been integral to studying the relationship between genotype and phenotypic fitness. These include functional genomic library screening/selection^4,5^, characterization of natural and synthetic cellular systems^6–8^, and directed evolution to either study evolutionary processes^9–11^ or evolve new biological function^12–15^. While identification of selected genotypes has dramatically improved with the advent of next generation sequencing, the ability to precisely and systematically control the conditions of selection has not improved in complimentary fashion, hampering understanding of how adaptive genotype is shaped by selective pressure^16^.

**Figure 1.**
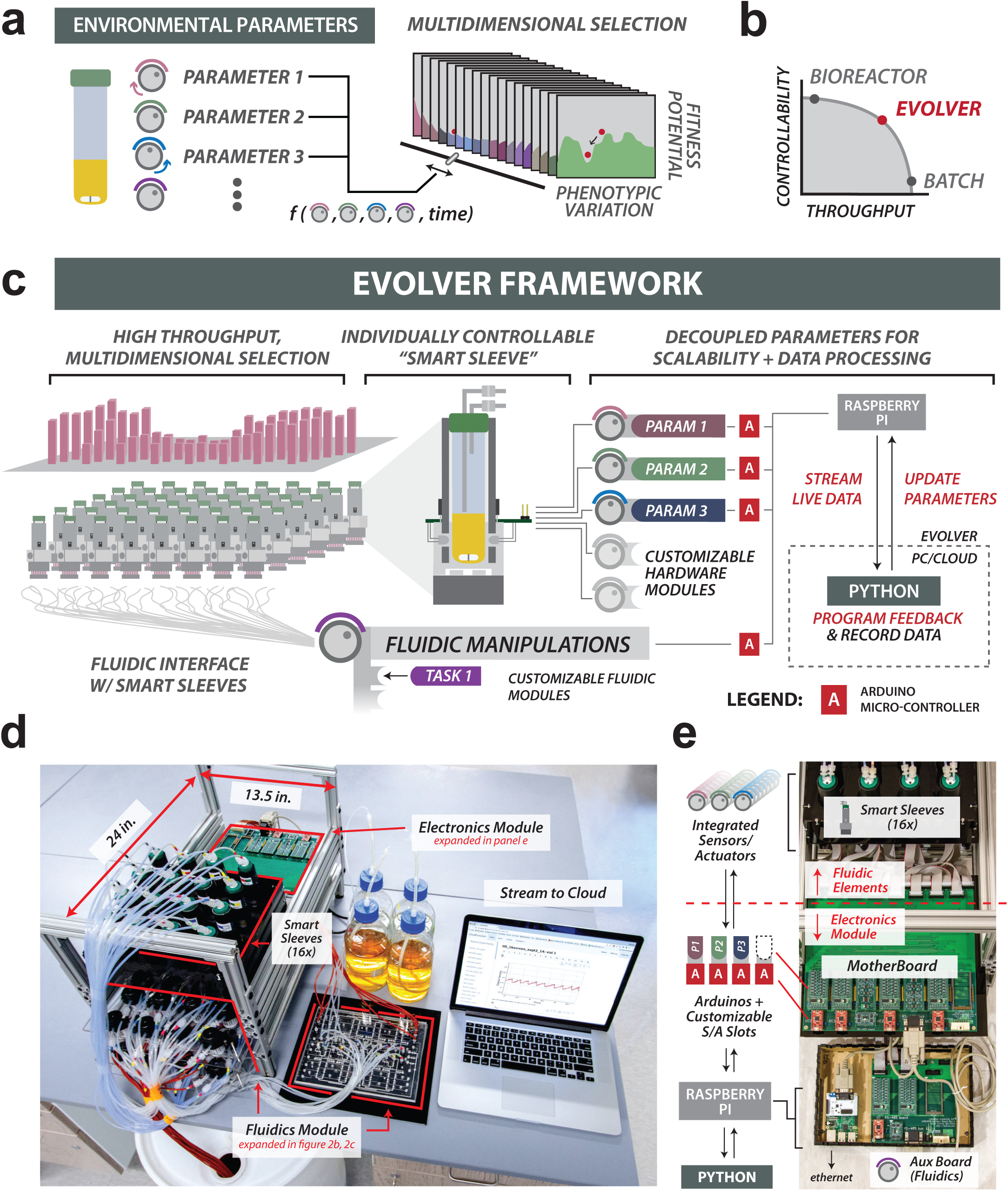
eVOLVER: an integrated framework for high-throughput, automated cell culture. **(a)** Understanding how cellular phenotypes arise from multidimensional selection gradients requires multi-parameter control of culture conditions. **(b)** Growth fitness experiments face a tradeoff between precision control of culture conditions and throughput. eVOLVER enables reliable scaling along both axes. **(c)** eVOLVER hardware, fluidic, and software modules. System design is modular and synergistic. Left: eVOLVER is designed to scale to high-throughput. Center Top: Smart Sleeve unit. Smart Sleeves integrate sensors and actuators needed to measure and control parameters of individual cultures. Center Bottom: eVOLVER fluidic manipulation system (peristaltic pumps or millifluidic devices) controls movement of media and culture within the system. Right: A modular, scalable hardware architecture interfaces with Smart Sleeve and fluidic modules to achieve individually addressable, real-time culture control. The hardware functions as a bidirectional relay, streaming live data (via Raspberry Pi) collected from each Smart Sleeve to the external computing infrastructure running control software (written in Python). This software records and processes data and returns commands to the hardware in order to update culture parameters. System customization can be achieved by swapping fluidic handling devices, adding new parameter control modules, or programming new feedback control routines between culture and software. **(d)** 16-culture eVOLVER base unit. Fluidics (media input, waste output) are physically separated from the electronics. The base unit can be cloned and parallelized to increase experimental throughput.**(e)** eVOLVER hardware architecture. Smart Sleeves communicate with electronics module via a motherboard. Control modules, which control single parameters across for all Smart Sleeves within a 16-culture unit, are composed of Arduino-connected control boards occupying motherboard S/A slots. Arduinos are programmed to interpret and respond to serial commands from the Raspberry Pi, which communicates with software run on a user’s computer or server.

An intrinsic challenge in designing growth selection experiments lies in balancing the tradeoff between control and throughput (**Fig. 1b**); while batch cultures maintained by serial passage permit parallel testing of many strains or cultures conditions, they are inherently discontinuous and offer limited temporal control over culture conditions^17^. Conversely, automated cell growth systems can maintain constant growth rates under precisely defined conditions, but are difficult to parallelize due to cost, space-inefficiency, and design complexity^16–18^. A recent convergence of several open-source technologies—inexpensive additive manufacturing, do-it-yourself (DIY) software/hardware interface, and cloud computing— has enabled the in-lab fabrication of new custom laboratory platforms^19–22^. Various examples in automated cell culture include devices designed to perform automated dilution routines for exploring antibiotic resistance acquisition^23^, or that implement design features such as real-time monitoring of bulk fluorescence^7^, light-based feedback control of synthetic gene circuits^24^, and chemostat parallelization^25^. Unfortunately, the single-purpose, ad hoc design of these systems limits their scalability and restricts their reconfiguration for other experimental purposes.

Here we present eVOLVER, a multi-objective, DIY platform that gives users complete freedom to define the parameters of automated culture growth experiments (e.g. temperature, culture density, media composition, etc.), and inexpensively scale them to an arbitrary size. The system is constructed using highly modular, open-source wetware, hardware, electronics and web-based software that can be rapidly reconfigured for virtually any type of automated growth experiment. eVOLVER can continuously control and monitor up to hundreds of individual cultures, collecting, assessing, and storing experimental data in real-time, for experiments of arbitrary timescale. The system permits facile programming of algorithmic culture ‘routines’, whereby live feedbacks between the growing culture and the system couple the status of a culture (e.g. high optical density (OD)) to its automated manipulation (e.g. dilution with fresh media). By combining this programmability with arbitrary throughput scaling, the system can be used for fine resolution exploration of fitness landscapes, or determination of phenotypic distribution along multidimensional environmental selection gradients.

We demonstrate the broad applicability of eVOLVER by configuring it to perform diverse growth and selection experiments. First, we conduct high-throughput experimental evolution on yeast populations using multi-dimensional selection criteria, scanning culture density space at fine resolution to assess adaptive outcomes. Next, by performing growth selection on a yeast knockout (YKO) library under temporally variable temperature stress regimes, we show that eVOLVER can be used to systematically explore the relationship between environmental fluctuations and adaptive phenotype. Finally, by integrating millifluidic multiplexing modules, we demonstrate that eVOLVER can carry out complex fluidic manipulations, dramatically extending the scope and range of possible growth and selection experiments.

## RESULTS

### eVOLVER: A Scalable Automated Cell Culture Framework

Most automated culture systems are designed to address specialized needs and are thus limited to active control over one or more predetermined culture parameters. We designed eVOLVER so that it could be flexibly configured to measure and control an arbitrary, user-defined set of parameters. In order to accomplish this, the system’s hardware design emphasizes rapid, cost-effective scaling and customization, as well as accommodation of future technological advancements (**Fig. 1c**). eVOLVER hardware includes the following three modules (**Fig. 1d**): (1) customizable Smart Sleeves, which house and interface with individual culture vessels, (2) a fluidic module, which controls movement of liquid in and out for each culture vessel, and (3) a plug-and-play hardware infrastructure that simplifies high-volume bidirectional data flow by decoupling each parameter into individual microcontrollers (**Fig. 1e**). A detailed description of eVOLVER design and construction can be found in **Supplementary Information 1** (resources also available online at **fynchbio.com**).

The Smart Sleeve is a manufacturable unit that mediates monitoring and control of growing cultures (**Fig. 2a**). Each sleeve is composed of a machined aluminum tube (for temperature control), printed circuit board (PCB) mounted sensors, actuators, and other electronic components, all attached to a custom 3D printed mount. As designed, the Smart Sleeve is one of the principal embodiments of eVOLVER versatility; it can be inexpensively mass-produced for high-throughput experiments, or reconfigured to meet custom experimental needs, such as larger culture volumes, pH or oxygen sensors for bioprocess applications, or light emitting diodes (LEDs) for optogenetic studies. In this work, we implement a particular Smart Sleeve configuration that accommodates 40 mL glass vials, and is configured to control three experimental parameters: stirring, temperature, and culture density (**Figs. 2a, S4, S5, S7**).

**Figure 2.**
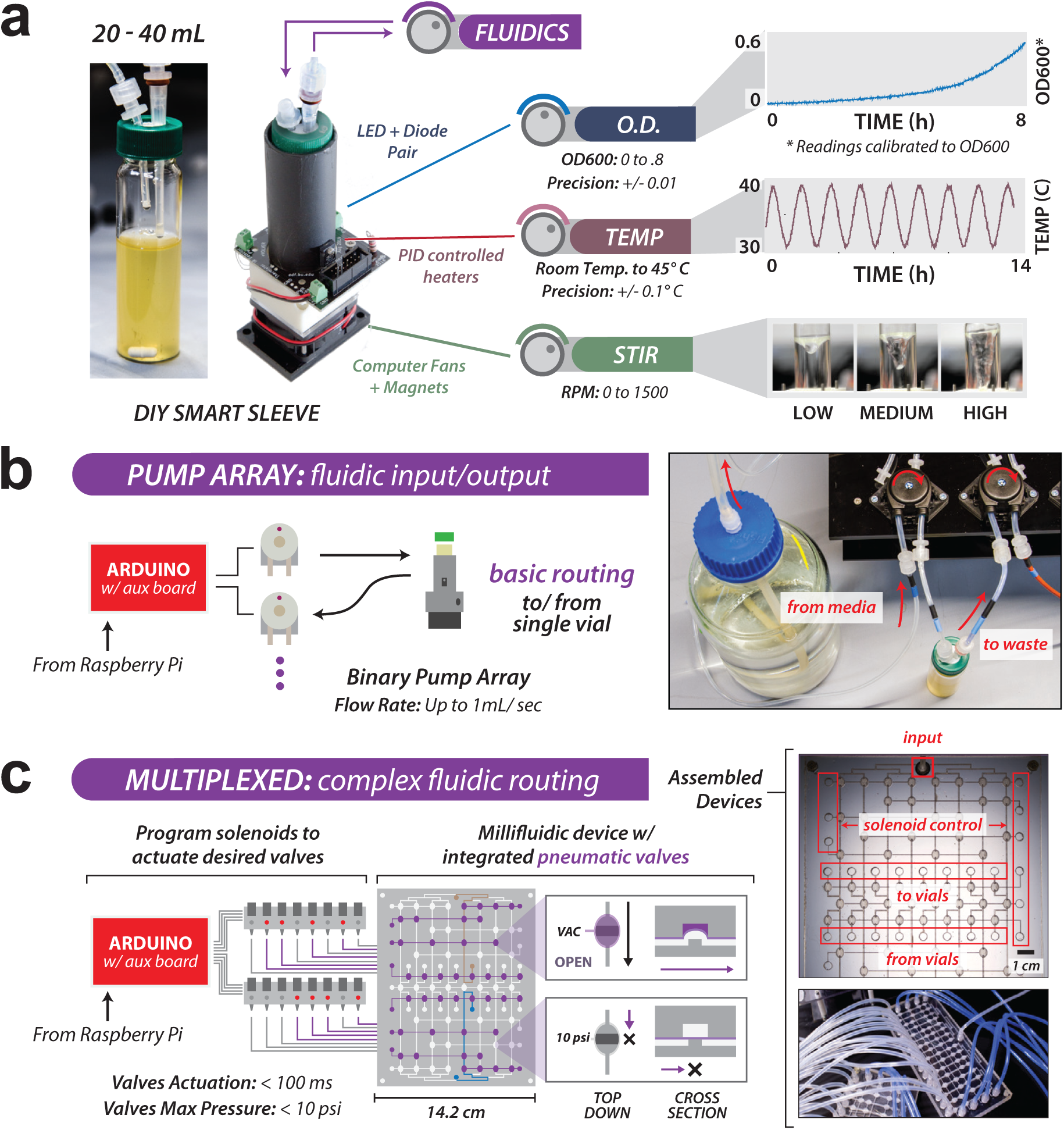
Design and performance of eVOLVER modules. **(a)** Generalizable configuration of Smart Sleeves for continuous culture; control of fluidic input/output, optical density, temperature, and stir rate. Left: Smart Sleeves are designed to accommodate 40 mL autoclavable borosilicate glass vials. Efflux straw length determines culture volume. Center: Smart Sleeve integrated electronic components. LED/photodiode sensor pairs perform OD_900_ readings. Thermistors and heaters attached to a machined aluminum tube maintain PID temperature control. Magnet-attached computer fans rotate stir bars inside the vials. Components are wired to a PCB and mounted on an inexpensive 3D printed chassis. Individual sleeves cost ∼$25 and can be assembled in ∼10 minutes. Right: Specifications of Smart Sleeve parameters: optical density, temperature, and stirring. Device measurement precision varies with experimental conditions (e.g. cell type, room temperature) but can be adjusted to achieve necessary precision and range (e.g. tuning temperature PID constants, or filtering OD measurements) (see **Supplementary Information 1**). Reported values are typical for experiments described in **Figs. 3-4**. Calibration may be performed as often as desired, though settings are largely invariant over thousands of hours of use. **(b)** “Basic” fluidic handling in eVOLVER utilizes pumps with fixed flow rates of ∼1 mL/sec and can be actuated with a precision of ∼100 ms. **(c)** Millifluidic multiplexing devices enable novel, customized liquid routing. Devices are fabricated by bonding a silicone membrane between two plastic layers with laser-etched flow channels. Integrated pneumatic valves actuate on the membrane to direct fluidic routing from media input to output ports (to or from vials).

eVOLVER’s fluidic module, which controls movement of media, culture, and liquid reagents within the system (**Figs. 2b-c, S9**), can be configured to two fluidic handling modes:(1) “basic” mode, which uses peristaltic pumping to control media influx and efflux for each culture^26^ (**Figs. 2b, S9b, S10**), and (2) a “complex” scheme in which multiplexed routing enables more sophisticated fluidic manipulations (**Figs. 2c, S9c**). User-actuated peristaltic pumps are robust, simple to use, and can be scaled to carry out media dilution routines for a large number of parallel continuous cultures^25,26^ (**Fig. S10**). However, they scale poorly to more sophisticated routing where the number of required connections and control elements expands non-linearly with the number of cultures. Our “complex” fluidic solution uses the principle of large-scale integration (LSI) to overcome this limitation. Originally developed for electronic devices, and then elegantly adopted in microfluidics^27,28^, LSI uses combinatorial multiplexing to expand the number of input-output paths per control channel^29,30^. Using LSI as inspiration, we created physically-compact millifluidic multiplexing devices by adhering a silicone rubber membrane between two clear sheets of laser-etched plastic, each patterned with desired channel geometries and aligned to form an intact device (**Figs. 2c, S11, S26, S27**). Devices can be designed on-the-fly to carry out custom fluidic protocols, including complex media dispensing routines, transfer of liquid between cultures, or periodic cleaning protocols to maintain sterility. Full treatment of multiplex device design and fabrication can be found in **Supplementary Information 1.5**, and a catalog of devices developed for this study can be found in **Supplementary Information 2.4**.

We developed a hardware and software infrastructure for eVOLVER that complements the scalability and configurability of the Smart Sleeve and fluidic modules (**Fig. 1c**). Our design is analogous to that of desktop computers, where a central organizing PCB (motherboard) is responsible for core functionalities (e.g. serial communication, signal routing), but also contains pluggable slots for boards that manage specialized features (e.g. graphics cards) (**Fig. 1e**). In eVOLVER, control modules—custom PCBs wired to Arduino microcontrollers—read and power Smart Sleeve-mounted components. During an experiment, individual control modules manage each culture parameter (e.g. temperature, stirring) (**Figs. S1, S2**). A motherboard distributes control of up to 4 independent parameters across a set of sixteen Smart Sleeves, comprising a single eVOLVER base unit (**Figs. 1d, S12**). Microcontrollers associated with each base unit are coordinated by a single Raspberry Pi, a small, low-cost, single-board computer that serves as a bidirectional relay to a user’s computer or a cloud server (**Figs. 1c, 1e, S3**). The modularity of this design facilitates repurposing and scaling; users can easily modify or augment eVOLVER’s experimental capability by connecting new control modules, while additional base-units can easily be cloned to achieve higher throughput (**Figs. 1d, S12**).

An important feature of eVOLVER’s design is its ability to leverage network connectivity to coordinate and run experiments over the internet. eVOLVER’s distributed hardware architecture enables efficient transmission of large packets of high-dimensional, real-time data (**Figs. 1c, 1e**) such that a single computer, located anywhere with an internet connection, can monitor hundreds of cultures in real time (**Fig. S3**). Python scripts running on the computer manage the acquired data and execute the control algorithms that define a selection scheme. For example, during a typical data acquisition/control protocol, a scripted routine may query the Raspberry Pi every 30 seconds for Smart Sleeve-acquired culture status data (e.g. temperature, optical density, etc.) (**Figs. 1c, 1e**). After being recorded, higher-order data (e.g. growth rate) are computed by an algorithm and used to update individual Smart Sleeves or fluidic channels with new settings (e.g. adjusted temperature or media dilution). By modifying a script, the user can change selection criteria between different modes of automated culture (**Fig. S37**), or specify a selection pressure surface through subtle iterations of the same algorithm across many Smart Sleeves.

eVOLVER is capable of robust long-term operation. The system configuration described in this paper is capable of running long-term (250+ h) experiments without electronics or software failure (**Fig. 3, S13**). Since most components have very long lifetime (>4000 h), hardware replacement is not required over the course of most experiments. Hardware calibration may be performed as frequently as desired, but we observed that settings remain essentially invariant over dozens of experiments (over 1700 h of operation) (**Figs. S4, S6, S8, S10**). eVOLVER is designed to be robust to catastrophic liquid spills, since Smart Sleeves and fluidic components are physically separated from their control hardware (**Fig. 1e**). Additionally, the system can be easily set up to avoid microbial contamination. In a control experiment, we found that the system remained sterile for the entirety of the experiment (∼10 days) while incubating uninoculated, antibiotic-free media alongside actively passaged *E. coli* cultures (**Fig. S13**).

**Figure 3.**
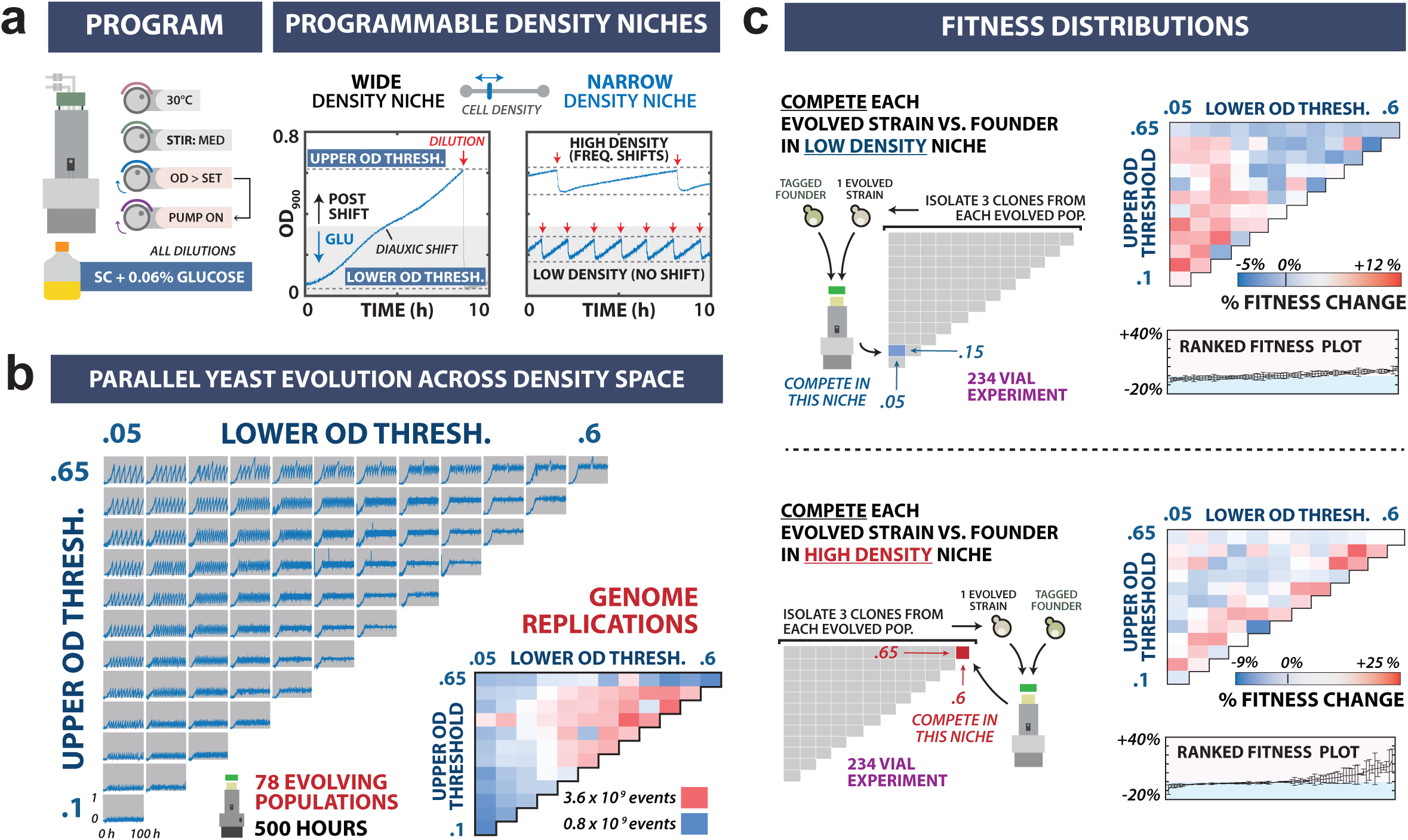
High-throughput experimental evolution across a multidimensional selection space. **(a)** Programming eVOLVER to maintain culture density selection routines during yeast evolution. Left: eVOLVER was configured to maintain cultures within defined density niches using a feedback between OD measurements and dilution events (turbidostat function). Right: Representative growth traces for yeast (*Saccharomyces cerevisiae* FL100) cultures growing under wide and narrow density niches. For each culture, the programmed OD window determines population size, and the consequent dilution rate and diauxic shift frequency. **(b)** Parallel evolution of 78 yeast populations in distinct density niches. Culture OD traces are shown for populations evolved for 500 h in density windows with varied lower (0.05-0.6) and upper (0.1-0.65) OD thresholds. Lower right: Heat map of the estimated genome replication events for the 78 populations. Values were calculated by multiplying average number of cells by the number of doublings, both estimated through segmentation of the OD trace. **(c)** Fitness distributions of evolved strains. Three clones from each evolved population were competed against the ancestral strain under low-density (OD 0.05-0.15, top) and high-density (OD 0.60-0.65, bottom) growth regimes. Right: Heat maps for mean fitness change relative to the ancestor (top) and ranked fitness with standard error bars representing competitive fitness for each clone (bottom).

### Conducting Experimental Evolution Across a Multidimensional Selection Space

In order to showcase eVOLVER’s ability to conduct long-term continuous culture laboratory evolution experiments at high throughput, we explored the relationship between culture density and fitness in evolving yeast populations. To accomplish this, we configured eVOLVER to function as a turbidostat, maintaining culture density within a constant, defined window bounded by lower and upper OD thresholds (OD_lower threshold_ – OD_upper threshold_). Using continuously recorded OD data (**Figs. S7, S8**), a routine activates dilution when the upper threshold is exceeded, and continues to fire until the lower threshold is reached (**Fig. 3a**).Population growth rates can be calculated in real-time by segmenting and fitting the OD trace between dilution events. We varied the upper and lower OD thresholds across 78 yeast populations grown in parallel, thereby defining a two-dimensional selection space based on minimum and maximum culture density (**Fig. 3b**) (see **Methods, Supplementary Information 2.2**). Cultures were grown in glucose-limited media for 500 h (40-280 generations depending on the culture), resulting in diauxic shifts in cultures with OD windows that exceed ∼0.35 (**Fig. S14**). The frequency with which the culture experiences the shift is a function of density-dependent selection prescribed to individual cultures (**Fig. 3b, S14**). Recorded OD measurements allowed us to generate maps of population and evolution parameters across the selection space, including population growth rates (**Fig. S15**) and average genome replication events, a function of OD and growth rate (**Fig. 3b, S16**).

We then used eVOLVER to perform fitness measurements on isolates from each of the evolved populations by competing them against a fluorescently-labelled ancestor strain under low- and high-density continuous growth regimes (468 total cultures), assaying for population ratio over time using flow cytometry^31^ (see **Methods**). The fitness distributions generated for low- and high-density regimes were distinct (**Fig 3c**). K-means analysis of the fitness phenotypes yielded three distinct groups: low-density specialists, high-density specialists, and the remainder exhibiting low fitness in both measured niches (**Fig. S17**). In general, low-density specialists evolved in niches with small lower OD thresholds. In contrast, high-density specialists were derived from cultures maintained in narrow, high-OD windows, rather than simply niches with high OD thresholds (**Fig. 3c, S17**). Our results demonstrate that in applying multidimensional selection pressures, eVOLVER can be used to map pressures that drive adaptation to specific regions of the selection space.

### Growth Selection Under Temporally Varying Selection Regimes

There is growing interest in understanding how an environment’s temporal features shape organismal adaptation^32^. A fluctuating environment may yield adaptations distinct from those selected under monotonic pressure^33,34^. eVOLVER can be used to systematically vary temporal features of a selective pressure while holding other culture conditions constant. To demonstrate this, we performed growth selection experiments on a pooled YKO library^4,5,35,36^ (5,149 unique members) under conditions in which a single environmental variable— temperature—was temporally varied. A two-dimensional selection space was programmed by varying the magnitude and period of square wave temperature oscillations (**Figs. 4a, S6**).Cultures were maintained for 6 days in turbidostat mode, and samples collected every 48 h (see **Methods, Supplementary Information 2.3**). At the conclusion of the experiment, next-generation sequencing was performed on each selected population to determine library member frequency^37,38^, which was used to calculate the fitness of each member^31^ (see **Methods, Supplementary Information 2.3**) (**Figs. S18-S20**).

**Figure 4.**
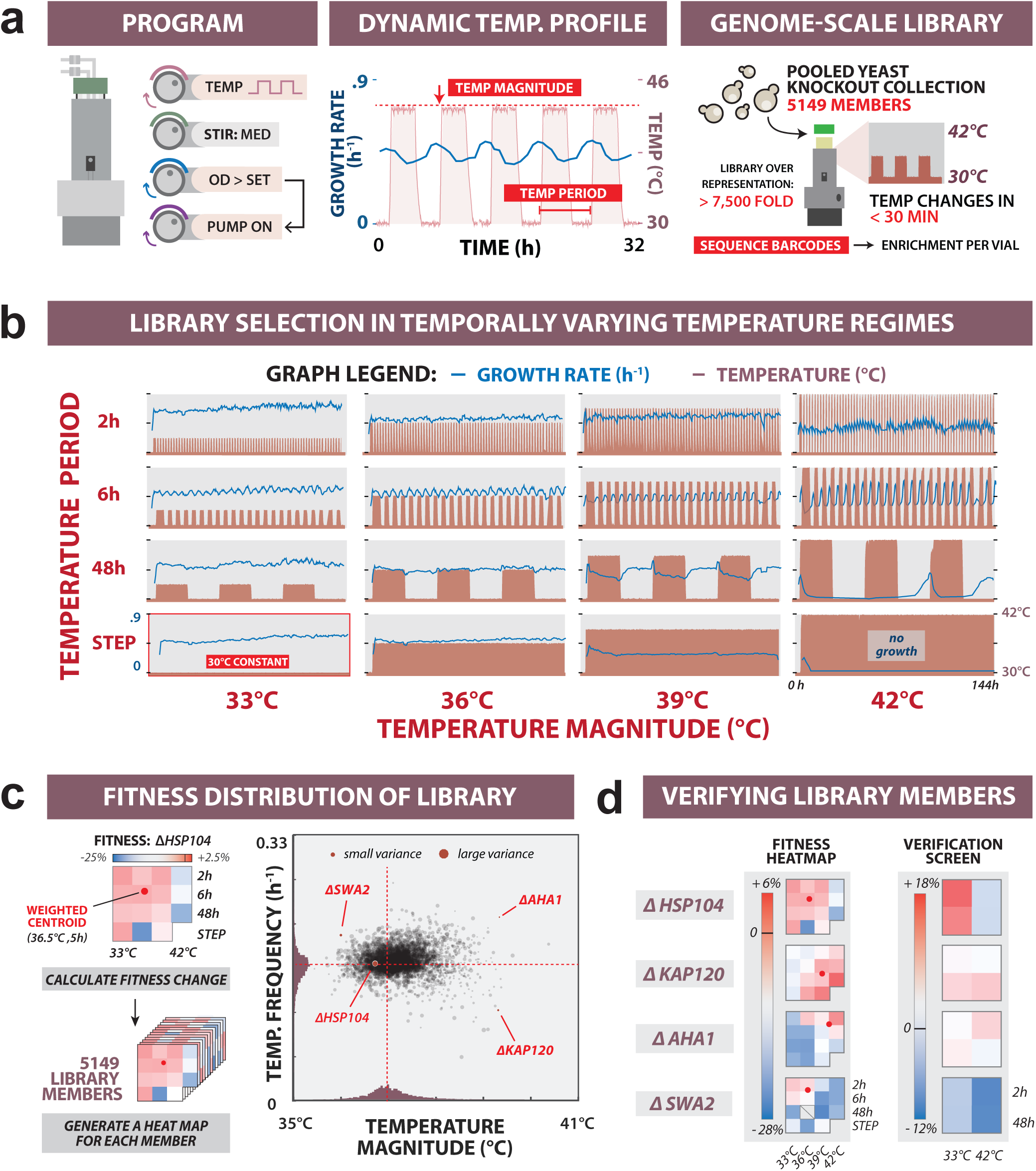
Genome scale library fitness under temporally varying selection pressure. **(a)** Programming temporally varying temperature regimes. Left: eVOLVER configuration for conducting turbidostat experiments (OD window: 0.15-0.2) under fluctuating temperature stress. Middle: Snapshot of temperature waveform (red) alternating between 30°C and 39°C on a 6 h period, and corresponding culture growth rate (blue). Right: Parallel cultures of pooled YKO collection were grown. Selection-based enrichment of library members was quantified at various timepoints using next-generation sequencing. **(b)** Full set of dynamic temperature regimes.Temperature magnitudes (33°C, 36°C, 39°C or 42°C) were varied against periods (2h, 6h, or 48h, or a constant step), and run against a 30°C control culture. Recorded temperature (red) is plotted with culture growth rates calculated between dilutions (blue). **(c)** Mapping fitness of library members to dynamic selection space. Left: For each library member, fitness heat maps were generated in each selection regime, and used to calculate weighted fitness centroids within temperature magnitude/frequency coordinate space. Right: Scatter plot of fitness centroids for the full library. **(d)** Validation of library selection. Four strains with distinct profiles were chosen for verification and competed against a neutral control strain (Δ*HO*) under four different temporal selection regimes. Population ratios were measured using quantitative PCR.

We computed a 2D weighted centroid for each library member by transforming its measured fitness into coordinates of temperature magnitude and frequency, allowing us to compare the library phenotypes across the 2D temporal selection space (**Supplementary Information 2.3**) (**Fig. 4c**). Plotting fitness centroids for the high-performing strains in each of the 16 conditions confirmed that the centroids cluster in the corresponding region of magnitude/frequency space (**Fig. S21**). Additionally, we picked four library members from different regions of the distribution (Δ*HSP104*, Δ*KAP120*, Δ*AHA1*, and Δ*SWA2*) for experimental validation of our library selection. We competed these strains against a neutral control strain (Δ*HO*) across four temperature profiles, observing fitness profiles that agree with the library selection results (**Fig. 4d**).

Library members with significant fitness centroid shifts along the magnitude or frequency axes were identified (**Fig. S22**), including several chaperone and chaperone cofactor genes, which are known to play a role in thermal stress response^39^. Saccharomyces Genome Database (SGD)^40^ phenotype annotations were then used to identify sets of similarly annotated genes with fitness centroids significantly shifted from that of the population mean (**Fig. S23**). Next, by applying cross-correlation and principle component analysis to compare each condition, we observed three distinct groups of conditions with correlated effects on the library: two high-temperature groups corresponding to (1) high- and (2) low-frequency, and (3) a mild temperature group (**Figs. S24, S25**). We identified gene ontology (GO) term annotations that are linked to fitness defects in one or more of these groups (**Fig. S25**). As expected, we observed that functions directly tied to growth rate (e.g. mitochondrial function, ribosome biogenesis) significantly altered fitness at mild temperature increases. Interestingly, ribosome components and processing factors also showed high-frequency sensitivity at high temperatures, suggesting a potential role for ribosome biogenesis in transitions in and out of stress. We further interrogated potential sources of frequency-dependence. We found that the high- and low-frequency groups were characterized by annotations associated with cell cycle checkpoints (e.g. DNA damage response, organelle fission), which temporally regulate cellular processes and thus might be expected to affect cellular response to fluctuating stresses at different frequencies.

### Enabling Complex Automated Culture Routines with Fluidic Multiplexing Devices

In order to demonstrate the potential of the fluidic multiplexing framework to automate movement of reagents and cells in eVOLVER, we designed devices for three experimental applications: (1) dynamic media mixing during continuous culture to track yeast response to ratios of sugars (**Supplementary Information 2.5**); (2) preventing bacterial biofilm formation via automated passaging (**Supplementary Information 2.6**); and (3) programming sexual reproduction between adapting yeast populations (**Supplementary Information 2.7**).

We showed that fluidic multiplexing could be used to manage media composition for multiple cultures maintained by eVOLVER by constructing an 8-channel media selector device that dynamically draws media from multiple input sources and dispenses a defined mixture to a culture of choice (**Supplementary Information 2.4**) (**Figs. 2c, S26, S27**). Galactose utilization in yeast is regulated by the ratiometric sensing of available galactose and glucose^41^. We maintained a yeast strain harboring a galactose-inducible fluorescent reporter (*pGAL1*-mKate2) in continuous cultures featuring 16 different sugar compositions (**Fig. S28**). Dyed media, tracking relative sugar levels in each culture, confirmed that the device can dynamically mix and dispense in correct ratios for extended periods of time (**Fig. S28**). We used flow cytometry measurements to track galactose-induced population fractions, confirming maintenance of sugar-dependent gene induction (**Fig. 5a**).

**Figure 5.**
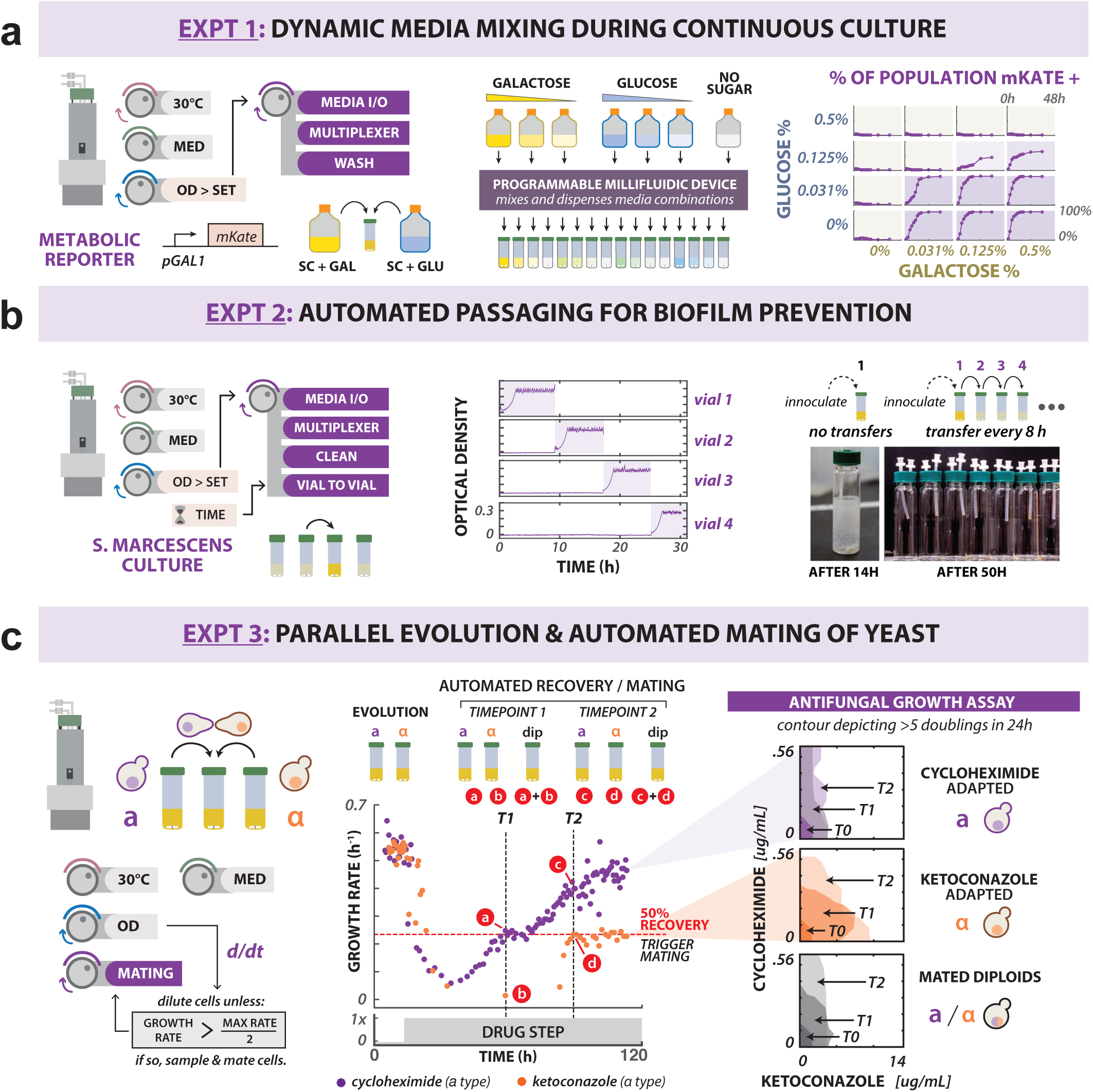
Integrated millifluidic devices enable scaling of complex fluidic manipulation. **(a)** Demonstrating dynamic media mixing in continuous culture. Left: eVOLVER program for maintaining cells in turbidostat mode using millifluidic device to mix and dispense appropriate dilution volumes. A yeast galactose-inducible reporter (*pGAL1*-mKate2) was used to validate the device by maintaining cultures in turbidostat mode at different ratios of glucose and galactose. Center: Any combination of seven media inputs can be mixed and dispensed into any of the 16 culture vessels. Right: Reporter induction (by population percentage) for 16 cultures containing different glucose:galactose ratios, as measured by flow cytometry. **(b)** Preventing biofilm formation with automated vial-to-vial transfers. Left: A millifluidic device can enable inter-culture transfers between any of the 16 cultures. Center: *Serratia marcescens* cultures were maintained in turbidostat mode, with culture transfer events triggered every 8 h. Right: A culture maintained in a single vessel forms a thick biofilm after 14 h, while automated transfer prevents visible biofilm formation. **(c)** Using millifluidic devices to automate yeast mating. Left: Haploid strains containing opposite mating types are maintained as turbidostat cultures under antifungal selection. Vial-to-vial transfers are triggered by growth rate feedback control, used to sample haploids and form diploids within the device using an automated mating protocol. Center: Growth rate of haploid cells evolved under cyclohexamide (CHX, 0.2 ug/mL, purple) or ketoconazole (KETO, 6 ug/mL, orange) selection was monitored continuously following drug exposure. Once growth rates of either drug-evolved culture equals 50% of the wild-type growth rate under no selection, automated mating and transfer is carried out. This was performed at two timepoints: t_1_=68.7 h and t_2_ = 98.1 h. Right: Antifungal resistance was assayed for recovered haploid and diploid populations. Contours correspond to an antifungal concentration range in which at least five generations of growth were observed in 24 h (on average).

The utility of the millifluidic system for mediating liquid transfer between cultures was demonstrated using a device that overcomes biofilm formation during long-term continuous growth experiments (**Fig. S27,S30).** *Serratia marcescens*^42^, a bacterial species that readily forms biofilm on glass surfaces, was grown in turbidostat mode. Using a vial-to-vial transfer device, inoculation of a fresh vial was performed at 8 h intervals, followed by automated sterilization of culture-exposed fluidic paths (**Fig. 5b**). While thick biofilm deposition was observed after only 14 h in a culture grown continuously in a single vial (no transfers), no deposition was observed in vials maintained under the transfer routine (**Fig. 5b**). With daily replacement of used vials, four Smart Sleeves could be used to passage a culture, biofilm-free, for an indefinite period of time (**Fig. S31**).

Finally, we demonstrated that multiple functionalities can be combined to automate complex fluidic handling routines. We designed a fluidic routine for automated sexual reproduction of adapting yeast populations by integrating multiplexed media selection, vial-to-vial transfer, and post-transfer device cleaning (**Supplementary Information 2.4**) (**Figs. S26, S27, S30, S32**). This routine was used to conduct mating between separate, opposite mating-type haploid yeast evolving under different antifungal selections: cyclohexamide (CHX) and ketoconazole (KETO) (**Fig. 5c**). Rather than manually sampling at arbitrary timepoints, once haploid cultures reached user-defined growth rate milestones, automatic sampling and mating were carried out on the device **(Figs. S32, S33)**. A minimum inhibitory concentration (MIC) assays performed on both diploid and parental populations indicated that the haploid-evolved CHX resistance was transferred to diploids in a dominant manner (**Figs. 5c, S34, S35**). Conversely, the general antifungal resistance that emerged under KETO selection appeared to be recessive. Further sequencing of KETO-evolved haploid strains revealed nonsense mutations in *ERG3* (**Fig. S36**), which has been shown to confer recessive resistance to azole antifungals^43^.

## DISCUSSION

eVOLVER achieves the goal of creating a standardizing framework for automated cell growth experiments. The system is designed from the bottom-up to be customizable and expandable; its DIY infrastructure provides researchers with the ability to design, easily build out, and share new experimental configurations and data. eVOLVER’s design utility is manifested at the component level: in Smart Sleeve design configurability, in the ability to specify custom liquid manipulation routines using the fluidic system, and in the modularity and composability of the hardware and software systems (**Figs. 1-2**). As a consequence, with straightforward modification, eVOLVER can be reconfigured to conduct any of the recently reported continuous growth studies (**Table S1**), or could replace tedious batch culture techniques used in a number of recent experimental evolution studies^9,11,32,44^. Additionally, new hardware components can easily be incorporated into the platform. For example, integration with emerging open-source pipetting robots would automate culture sampling, unlocking downstream fluorescence-activated cell sorting (FACS) for assaying gene expression, or droplet microfluidics for single-cell studies. The present configuration is designed for well-mixed liquid cultures, but the eVOLVER control framework could be adopted for the coordination of multiple arrayed sensors to capture spatial distributions in static liquid cultures or phototrophic cultures. Finally, while eVOLVER is well suited for hardy, fast-growing suspension cultures of microbes, additional attention to sterility and removal of residual cleaning agents would be needed for sensitive mammalian lines, and bead/matrix systems may be needed for adherent cells.

The results reported in this study establish eVOLVER as a scalable framework for realizing large-scale, multidimensional selection experiments to study, characterize, and evolve biological systems. The system’s configurability enables precise specification of culture environment on an individual culture basis. By systematically co-varying parameters, eVOLVER can be used to investigate cellular fitness along multidimensional environmental gradients, potentially allowing for experimental decoupling of overlapping selection pressures. The ability to arbitrarily program feedback control between culture conditions and fluidic functions allows the user to algorithmically define highly specialized environmental niches.

We demonstrated the breadth of eVOLVER’s experimental versatility in a series of showcase experiments. First, we used eVOLVER to conduct an experimental evolution study (**Fig. 3**). There has been longstanding interest in the interplay between environmental carrying capacity, growth rate, and population size^45–47^. In our study, we evolved yeast across 78 different culture density windows. We then generated fitness distributions by testing fitness of evolved clones in low- and high-density niches, identifying low- and high-density specialists. Interestingly, high-density specialists were most often derived from evolution in narrow OD windows. Since the prescribed culture density windows are related to the frequency of diauxic shift at limiting glucose concentrations^48^, it is interesting to speculate that these strains selected for metabolic programs that facilitate rapid metabolic shifts. Critically, further work is needed to confirm that the differences observed in the fitness distributions are indeed significant. First, a baseline fitness distribution generated from a large number of replicate evolutions could be used to rule out the possibility that stochastic events dominate the observed fitness differences.Additionally, comparing the fitness of whole evolved populations in each condition could help isolate true adaptation from variation observed due to clonal differences. Finally, assaying fitness in additional niches is required to determine how well fitness distributions correlate with the assayed niche, as well as to ascertain the existence of generalists. Nevertheless, this experiment demonstrates how eVOLVER’s high-throughput capabilities can be used to uncover subtly varied adaptations that likely would have been obscured by either a batch culture or a lower-resolution automated approach. Furthermore, eVOLVER enables rich phenotypic profiles to be constructed from individual culture histories using data collected during long term experiments (e.g. growth rate, genome replications).

In a second experiment, we performed growth fitness experiments on a YKO library under systematically varied temperature fluctuations, demonstrating that eVOLVER could be used to extract unique fitness information across a temporally varied selection surface (**Fig. 4**). In a similar approach—using sub-lethal stress administered under temporally diverse selection conditions—selection experiments could be conducted for other types of libraries at a resolution beyond what is available for standard growth selection schemes^4,37,38^. Using this approach, new algorithms with additional parameters can be used to perform fitness and selection experiments investigating biological phenomenon associated with temporal adaptation, like bet-hedging, noise, and transcriptional feedback.

Accurate fluidic manipulation is a critical feature of continuous culture automation, but past approaches to fluidic routing have imposed experimental limitations. Inspired by electronic and microfluidic technologies^27^, we addressed this issue by developing a novel millifluidic control paradigm for programmatic routing of fluids during continuous culture. Our devices expand the repertoire of cellular manipulations available to eVOLVER while packaging functionality into a compact physical footprint. To highlight the utility and robustness of the devices, we performed three experimental demonstrations: sophisticated fluidic mixing and dispensation, vial-to-vial transfers, and integration of multiple devices for more complex culture routines (**Fig. 5**). These experiments illustrate the potential of custom millifluidics to create sophisticated algorithmic selection routines. As we demonstrate, eVOLVER can be configured to culture separate strains, simultaneously intermixing them to study community interactions.For microbial populations capable of exchanging genetic information, such an approach could be used to explore different paths by which resistance can spread, with implications for infectious disease treatment.

We foresee the eVOLVER platform as an enabling tool for several emerging fields of research. eVOLVER could play an important role in investigating the adaptive basis of social behavior in microbial consortia. For example, the system’s throughput, control, and fluidic capabilities could be leveraged to systematically test contributions of individual species to community fitness, potentially offering insight into how to construct ecologically stable communities from the bottom up^49^. In synthetic biology, designing regulatory circuits that minimize fitness cost to the host cell remains a major challenge^3^. Leveraging its ability to carefully monitor population fitness in high throughput, eVOLVER could be used to identify circuit design features that maximize evolutionary stability. This would be useful for applications requiring engineered cells to retain circuit function over many generations, and would be especially valuable for high-throughput testing of bioproduction strains prior to scaling-up to industrial bioreactors^50^. Relatedly, eVOLVER could be used to aid in the design, testing and optimization of synthetic microbial genomes^51,52^. Finally, we envision eVOLVER as enabling platform for building a community of users that can freely conceive, build, execute, and share experiments.

## METHODS

General methods and analysis used throughout the study are described here. A detailed description of eVOLVER device design can be found in **Supplementary Information 1**. For a more complete description of experiments depicted in **Figs. 3-5**, we refer the reader to **Supplementary Information 2**, which includes detailed methods, results, and data analysis.

### Strain Construction

Genotypes for yeast and bacterial strains used in this study are listed in **Table S2**. Plasmids were constructed using standard molecular biology techniques. Strains were generated using standard lithium acetate transformation. When appropriate, non-isogenic pooled population samples were given unique designations for clarity (**Table S2)**.

### Routine Cell Culture Techniques

Culture conditions varied according to the needs of particular experiments (see methods in **Supplementary Information**). Cultures used to seed eVOLVER experiments were prepared as follows: *Saccharomyces cerevisiae* (obtained from frozen stock or single colonies) was grown in 2 mL of YPD media (2% glucose) at 30°C in a shaking incubator (300 rpm) for at least 36 h. For routine overnight culture of *Escherichia coli* or *Serratia marcescens*, cells obtained from frozen stock were used to inoculate 2 mL of LB Miller broth grown at 37°C for 12 h in a shaking incubator (300 rpm).

### Flow Cytometry

Flow cytometry was used to measure single-cell fluorescence throughout the study. Prior to measurement, 200 uL of yeast culture (see methods for experiment-specific growth conditions) was diluted with 100 uL of filter-sterilized PBS supplemented with cyclohexamide to a final concentration of 20 ug/mL, then incubated at 4°C in the dark for no less than 3 h to allow for fluorophore maturation. An Attune NxT Flow Cytometer (Invitrogen) equipped with an autosampler was used to acquire data. For a typical experiment, at least 10,000 events were acquired. Cells were analyzed using FlowJo (Treestar Software). Intact cells were gated using forward and side scatter, followed by gating on fluorescence channels (green and/or red, as appropriate) to determine the fractional distribution of each population.

### Fitness Calculations

Competitive fitness, in which a strain of interest is co-cultured in competition with a reference strain, was assayed in the same fashion throughout the study. The ratio of the two strains was determined at multiple timepoints – generally at the beginning and end of an experiment – either by flow cytometry or qPCR. Fitness values, *F*, were calculated using the following equation^31^:

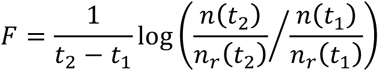

where *t* is number of generations, and *n* and *n*_*r*_ are cell counts for the strains of interest and reference strain, respectively.

### Setup Procedure for eVOLVER Experiments

Prior to each eVOLVER experiment, 40 mL borosilicate glass vials outfitted with a stir bar and capped with an influx port and an efflux port were sterilized by autoclave. Media and waste lines were sterilized by pumping 10% bleach (20 mL), followed by 70% ethanol (20 mL). Lines were cleared by pumping air for 20 s, followed by media (20 mL). Media and waste lines were then attached to the influx and efflux ports of each vial, and each vial filled by pumping 25 mL of the appropriate media through the influx port. At this point, Python control code was initiated, triggering a blank media measurement, and activating Smart sleeve heating elements. Prior to seeding with cells, fresh media was incubated in the device for a length of time sufficient for the first 15 optical density recordings to be taken (∼2.5 min) and for media to reach the programmed temperature.

### Extracting Yeast Genomic DNA

To extract genomic DNA, ∼2 x 10^6^ yeast cells (roughly 30 µL of overnight culture) were pelleted by centrifugation (5 min, 1000 rcf). Supernatant was removed, and pellets were resuspended in 30 µL 0.2% sodium dodecyl sulfate (SDS), followed by vortexing for 15 s. Suspensions were transferred to PCR tubes and heated in a thermal cycler (37°C) for 5 min, followed by 98°C for 5 min before cooling to 4°C. Extracts were diluted with H_2_O to a final volume of 75 µL prior to being used as PCR template. Primers used in this study are listed in **Table S2**.

### Library Preparation and Barcode Sequencing

Libraries for next-generation sequencing were prepared by PCR-amplification of genomic DNA (LightCycler 480 Instrument II, Roche), purification (Zymo Research). Primers used for indexing (culture, timepoint) and sequencing adapters were added via PCR. NextSeq sequencing (Harvard Biopolymers Facility) was used to sequence the culture index, the timepoint index, and a 55 bp single end read of the barcode construct. PhiX was added at 50% to increase sequence diversity. Alignment was performed using custom code harnessing MATLAB’s Bioinformatics Toolbox and Boston University’s parallel computing cluster. Alignment scores were calculated using the Smith-Waterman algorithm (swalign function) and assigned based on best score above a minimum threshold. In total, we assigned more than 244 million reads to 5149 unique library members to track frequency across the 4 time points for each of the 16 conditions. A more detailed description of the library preparation and sequencing can be found in **Supplementary Information 2.3.**

## Data Availability

Data supporting these studies are available upon reasonable request.

## ACKNOWLEDGEMENTS

We are grateful to B. Stafford for his assistance in design architecture of the system. We thank H. Khalil, A. Soltanianzadeh, A. Sun, S. Pipe, and A. Cavale for help on construction of the system. We are indebted to the Electronics Design Facility (EDF) and Engineering Product Innovation Center (EPIC) at Boston University for their services. We also thank D. Segrè, J. Ngo, J. Tytell, W. Wong, and members of the Khalil Lab for insightful comments on the manuscript. This work was supported by a NSF CAREER Award (MCB-1350949 to A.S.K.) and a DARPA grant (HR0011-15-C-0091 to A.S.K.). A.S.K. also acknowledges funding from the NIH Director’s New Innovator Award (1DP2AI131083-01), DARPA Young Faculty Award (D16AP00142), and NSF Expeditions in Computing (CCF-1522074).

## CONTRIBUTIONS

B.G.W, C.J.B, and A.S.K. conceived the study. B.G.W. developed the system with guidance and input from all authors. B.G.W and C.P.M. performed and analyzed experiments. S.K. generated reagents. C.J.B. and A.S.K. oversaw the study. All authors wrote the paper.

